# Degree-day-based model to predict egg hatching of *Philaenus spumarius* (Hemiptera: Aphrophoridae), the main vector of *Xylella fastidiosa* in Europe

**DOI:** 10.1101/2022.11.10.515963

**Authors:** Clara Lago, Alex Gimenez-Romero, Marina Morente, Manuel A. Matías, Aránzazu Moreno, Alberto Fereres

**Author notes:** Corresponding author: Alberto Fereres. ICA-CSIC C/Serrano 115b. 28006, Madrid, Madrid. Spain.

## Abstract

*Philaenus spumarius* L., the main vector of *Xylella fastidiosa* (Wells) in Europe, is a univoltine species that overwinters in the egg stage, and its nymphs emerge in late winter or spring. Predicting the time of egg hatching is essential for determining the precise times for deploying control strategies against insect pests. Here, we monitored *P. spumarius* eggs from oviposition to egg hatching together with the daily temperatures and relative humidities at four field locations that were located at different altitudes in central Spain. The collected data were used to build a growing degree day (GDD) model to forecast egg hatching in the Iberian Peninsula. Furthermore, the model was validated with field observations that were conducted in Spain. The model was then used as a decision-support tool to calculate the optimum timing for applying control actions against *P. spumarius*. Our results suggest that controlling nymphs at two different dates would target the highest percentages of nymphal populations present in the field. Our model represents a first step for predicting the emergence of nymphs and adopting timely control actions against *P. spumarius*. These actions could limit disease spread in areas where *X. fastidiosa* is present.

## INTRODUCTION

Predicting epidemics can be challenging, especially after the emergence of a new pathogen in a novel ecosystem. *Xylella fastidiosa* Wells (1987) is a vector-borne plant pathogenic bacterium that is native to the Americas that has been recently introduced in Europe and the Mediterranean basin and detected so far in Italy, France, Spain, Portugal, Israel and Iran (EPPO 2019, EFSA 2022). This bacterium is the aetiological agent of several diseases that affect important crops, including Pierce Disease (PD) on grapevines, Almond Leaf Scorch Disease (ALSD) on almond trees, Citrus Variegated Chlorosis (CVC) on citrus and Olive Quick Decline Syndrome (OQDS) on olive. The meadow spittlebug, *Philaenus spumarius* L. (1758) (Hemiptera: Aphrophoridae), is the only epidemiologically relevant vector of *X. fastidiosa* in Europe (EFSA 2015, Cornara et al. 2019). Thus, a detailed understanding of its ecology and phenology is essential for developing an accurate forecasting model for controlling *P. spumarius*.

*Philaenus spumarius* is a univoltine species, with a single generation per year, which has an ovarian parapause and a winter diapause in the egg stage (Morente, Cornara, Plaza, et al. 2018, Antonatos et al. 2019, Bodino et al. 2020). Eggs overwinter until hatching occurs in early spring. After egg hatching, the pre-imago (nymph) passes through five instars and nymphal development takes about 5-6 weeks until they become adults (Weaver and King 1954, Whittaker 1965, Ossiannilsson 1981, Fielding et al. 1999). *Philaenus spumarius* adults are extremely polyphagous, while nymphs are more host-plant-specific compared to adults (Halkka et al. 1967, Ossiannilsson 1981). Both nymphs and adults spend most of their life cycle on ground cover herbaceous vegetation, where mating, oviposition and feeding occur. In Mediterranean climates, in late spring, when the ground vegetation dries out, spittlebug adults can migrate long distances (more than 2 km far) from the ground cover to trees and shrubs (Lago, Garzo, et al. 2021, Lago, Morente, et al. 2021, Casarin et al. 2022). Adults return in the fall to olive and other woody crops and females lay their eggs on plant debris on the soil until they naturally die during winter (Morente, Cornara, Plaza, et al. 2018, Dongiovanni et al. 2019, Antonatos et al. 2020). Eggs are laid in masses and covered by a cement substance and overwinter until hatching occurs in early spring (Weaver 1951, Whittaker 1965, Witsack 1973).

The study of the developmental times of ectotherms as a function of temperature, in particular insects, has a long history (Rebaudo and Rabhi 2018). Insects need specific accumulations of heat units (HU) to reach certain development stages, which are commonly defined by the growing degree days (GDD) (also referred to as degree-days, heat units or thermal units) (Herms 2004). Basically, the GDD is a measure of heat accumulation over time-based on insect development rates at temperatures between the lower and upper limits. Usually, the function that describes the temperature response exhibits a unimodal form, with minimum and maximum temperatures below or above which no development occurs. The minimum or base temperature (*T_base_*) at which insects begin to develop is known as the “lower developmental threshold” or “base temperature”, while the maximum temperature (*T_max_*) at which insects stop developing is called the “upper developmental threshold” or “cut-off” (Murray 2020). These minimum and maximum temperatures give rise to an optimal temperature (*T_opt_*) that is obtained by fitting a unimodal-shaped curve.

Degree-day models based on heat accumulation have commonly been used to support integrated pest management (IPM) programs for several insect pests. Diaz et al. 2007 established the temperature threshold for the development of *Nasonovia ribisnigri* Mosley (Hemiptera: Aphididae) by determining the developmental rates at constant temperatures. Likewise, in a study conducted by Kimberling and Miller 1988, eggs of the winter moth *Operophtera brumata* L. (Lepidoptera: Geometridae) were subjected to fixed temperatures to calculate their developmental threshold. Similar studies have improved the control measures against serious pests such as the spotted Lanternfly *Lycorma delicatula* White (Hemiptera: Fulgoridae) (Smyers et al. 2021) and citrus flatid planthopper *Metcalfa pruinosa* Say (Kim et al. 2020). Moreover, Singh et al. (2017) developed a GDD model to forecast the occurrence of *Aphis craccivorain* Koch (Hemiptera: Aphididae) in lucerne fields by using the heat accumulation values starting on a given date.

Regarding the meadow spittlebug, several authors have established correlations between *P. spumarius* phenology and temperatures. Bodino et al. (2019) calculated the days of development (DD) of the nymphal stages in the Apulia and Liguria regions of Italy, as a function of the number of hours in one year that was above a temperature threshold. The developmental thresholds were calculated from the data obtained by experiments performed at fixed temperatures. Beal et al. (2021) adapted the formula from Bodino et al. (2019) to study *P. spumarius* phenology in north coastal California. They obtained similar results, although slightly prolonged development for *P. spumarius* was observed in California compared to Italy. In contrast, Chmiel and Wilson (1979) estimated the upper and lower thresholds for the nymphal development of *P. spumarius* in America by using the lowest coefficient of variation (CV) method described by Arnold (1959). Moreover, they calculated the HU by using the method described by Sevacherian et al. (1977). Similarly, Zajac et al. (1989) fitted linear regression functions to study the nymphal development of *P. spumarius*. Furthermore, some authors have suggested that humidity plays a key role in the development of *P. spumarius* (Weaver and King 1954, Grant et al. 1998).

In many studies, a simple linear approximation was used to compute the GDD metric and used only the minimum temperature threshold (Johnson et al. 1998, Campbell and Hanula 2007, Bodino et al. 2019, Beal et al. 2021). Alternatively, complex nonlinear approximations have been used (Logan et al. 1976, Lactin et al. 1995, Briere et al. 1999). However, (Quinn 2017) concluded in his extensive critical review that for the majority of studies conducted thus far, more complex functions performed poorly relative to simpler ones. Moreover, degree-day models are usually based on laboratory assays that aim to experimentally determine the minimum, optimum and maximum development temperatures, and such assays typically involve fixed temperature-controlled experiments, (e.g. Diaz et al. 2007).

In the present study, we developed a degree-day model to forecast the egg hatching of *P. spumarius* by following the logical steps for model construction (Overton 1977): 1) Data collection: We obtained field data from independent experiments conducted on egg hatching of *P. spumarius* in their natural environments and on-site temperature recordings at specific locations to avoid the inherently noisy temperature time series that are measured in the field. 2) Model construction: Our model was based on a multilinear temperature response function with the minimum, optimal and maximum temperatures. 3) Model calibration: We calibrated the model and obtained the minimum, optimal and maximum temperatures that control egg development by using an optimization procedure to determine the parameters that provided the best fit to the available experimental data rather than by running laboratory observations at fixed temperatures. 4) Model validation: Our model was validated with systematic and independent survey data of newborn nymphs that were obtained by researchers who conducted field surveys of *P. spumarius* in different regions in Spain. 5) Model extension: The model was extended to predict egg hatching throughout the entire territory. Although our approach is based on relatively simple experimental measures, it provides accurate egg-hatching predictions in the Iberian Peninsula. Moreover, our GDD model was used as a decision support tool to determine the best timing for applying control measures to manage vector populations. This model could be used by farmers and farm practitioners to optimize control measures against *P. spumarius*, which is the main vector of *X. fastidiosa* in Europe.

## MATERIALS AND METHODS

### Insects and Plants

A total of 100 adult individuals of *P. spumarius* were collected at “Pinilla del Valle” (Madrid, Centre Spain) (coordinates: 40.611108, −4.263008) during spring-summer, 2020, and were maintained inside bug-dorm cages (0.5×0.5×0.5 m) on *Sonchus oleraceus* L. plants (4-5 leaf stage) in a net house with no environmental controls (Temperature °C: MEAN+SE=16+0.3, Max: 31.7, Min: 7.4; Humidity RH%: MEAN+SE=63.5+1.3, Max: 99.6; Min: 20.0) at the ICA-CSIC facilities (Madrid, Spain). The proportion of sexes was one female per two males to ensure mating (Morente, Cornara, Moreno, et al. 2018, Morente et al. 2021). To obtain egg masses for the assays, which were conducted in October 2020, dry pine needles were placed on the substrate below the *S. oleraceus* plants to facilitate oviposition (Morente, Cornara, Moreno, et al. 2018). The pine needles were checked once per week from 6-X-2020 to 4-XI-2020 to identify the egg masses (Figure 1A, 1B). The dates when egg masses were observed on the pine needles were recorded. A total of 262 egg masses from five different oviposition dates were used in the field assay. The oviposition periods were the days preceding 1) 8-X-2020, 2) 14-X-2020, 3) 22-X-2020, 4) 29-X-2020 and 5) 4-XI-2020.

**Figure 1:**
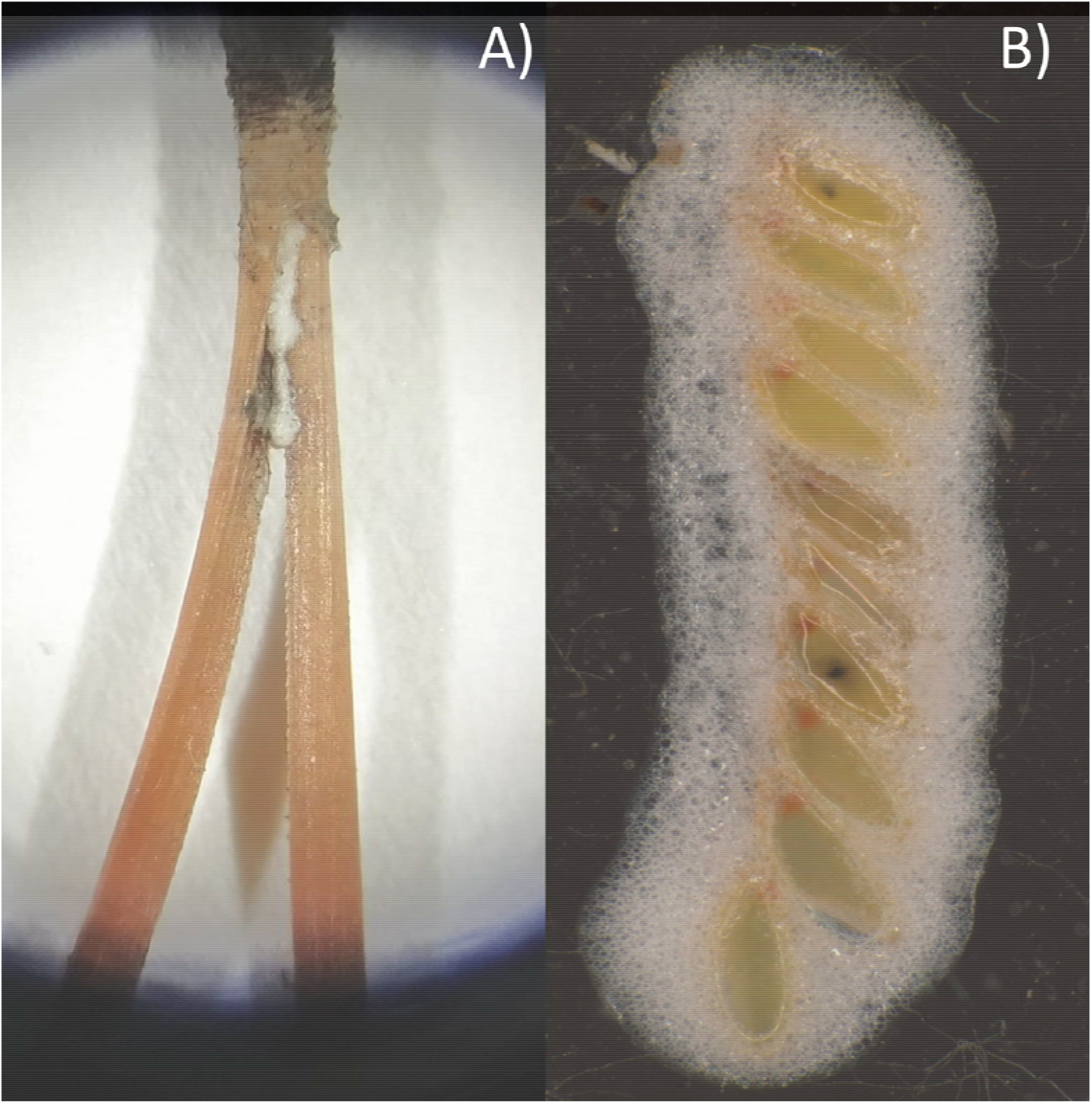
A) *Philaenus spumarius* newly laid egg masses on a dry pine needle. B) *Philaenus spumarius* egg mass.

### Data Collection: Monitoring egg hatching of *P. spumarius* under field conditions

The egg masses from each oviposition date that were obtained as described above were divided into four equal parts, and each part was transferred to a different field location on the day after the eggs were collected. The four locations used for our field study consisted of natural environments located in central Spain. These sites were selected at different altitudes to provide a gradient in climate conditions: 1) Alcalá de Henares (588 m) (coordinates: 40.521133, −3.290865); 2) Bustarviejo (1,222 m) (coordinates: 40.691827, −3.767162); 3) Mataelpino (1,086 m) (coordinates: 40.736439, −3.944309); and 4) Pedrezuela (880 m) (coordinates: 40.761003, −3.619030). At each field location, the egg masses were kept on an open 5.5-cm-diameter petri dish with small holes below to facilitate drainage (Supp. Figure S1). The petri dishes with egg masses were placed inside floorless mesh cages (50 cm height and 45 cm diameter) (Fyllen cloth basket 79 l, Inter IKEA Systems, Sweden). Two cages were placed at each field location: The cages were divided into four equal sectors by using cellulose acetate sheets. Since we had five oviposition dates, two and three plants were placed inside each of the two cages, respectively (Supp. Figure S2). Egg masses from a given oviposition date were placed in a single plant and sector of the cage to avoid mixing nymphs after emergence. A few weeks before the first nymphs were expected to emerge in the field (e.g., 25-II-2021), one *S. oleraceus* plant was transplanted inside each cage division where the egg masses were placed to feed the newborn nymphs after emergence (Supp. Figure S2). After transplanting, the egg masses were transferred from the petri dishes to the bare soil below the plants. In Mataelpino, the potted *S. oleraceus* plants inside the cages were watered weekly as opposed to the rainfed field locations. At this time, we checked each plant once a week to record egg hatching when newborn nymphs were observed. The newborn emerged nymphs were removed from the plants to facilitate further observations. All plants were inspected until no newborn nymphs were observed for two consecutive weeks.

Temperature and relative humidity (RH) were monitored hourly at each field location with data loggers (OM-EL-USB-2, Omega Engineering, INC, Norwalk, Connecticut, USA) during the entire duration of the experiments. One data logger not exposed to direct sunlight was placed inside each cage (Supp. Figure S1) at each field location to obtain the same temperature and RH data as that experienced by the egg masses. The data collected from the data loggers were downloaded every two weeks until the emergence of newborn nymphs. Thereafter, data were downloaded once a week. In the few cases where the data loggers failed to record weather data due to extreme weather conditions, the missing data were supplemented with external data collected from the nearest official meteorological station (Agencia Estatal de Meteorología de España, AEMET, Spain).

### Model Construction: GDD-based model

In the current study, we used a multilinear temperature response function with minimum or base (*T_base_*), optimal (*T_opt_*) and maximum (*T_max_*) temperatures to calculate the hourly contributions to the GDD. The use of a multilinear function is based on the principles of biochemical kinetics (i.e., on Arrhenius’ Law) (Eq. 1) and is explained in detail in Supp. Document S3.

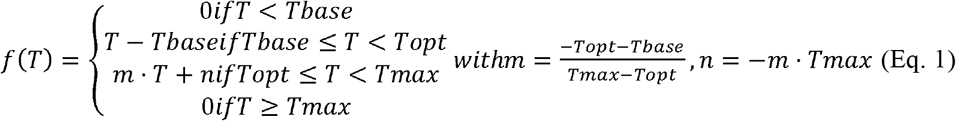

In our model, the accumulated GDD values are directly related to the cumulative probabilities of egg hatching using the cumulative density function of the Weibull distribution where k>0 and λ>0 are the shape and scale parameters of the Weibull distribution, respectively.

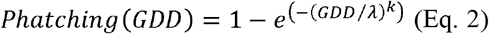

The starting date for GDD accumulation was the date when the winter diapause of *P. spumarius* ended. This date, in principle, should be determined by some metric of accumulation of cold temperatures (below a given threshold). However, the precise hours or days needed under a given temperature -cold requirement- and the precise length of the winter diapause of *P. spumarius* are unknown, Thus, we compared different arbitrarily fixed dates of diapause ending (January 1^st^, December 1^st^ and November 1^st^), and later validated the model with data obtained from field observations of newborn nymphs in the field from systematic surveys at different sites in Spain over the last six years (Supp. Table 4).

### Model Calibration

The goal of our model was to provide estimates of the cumulative hatching probabilities of *P. spumarius* eggs by using the temperature data provided by the field experiments described above transformed into a GDD metric. All data were treated independently of location. The experimental cumulative hatching probability at a given time was simply calculated as the number of nymphs that had already emerged at that given time over the total number of nymphs that had emerged by the end of the assay. Then, the cumulative GDD values between the end of diapause and the egg hatching date had to be calculated so that the cumulative hatching probabilities could be expressed as functions of the accumulated GDD values. In this way, we could fit Eq. (2) to the experimental data and obtain an effective model for predicting egg hatching. However, the relationship between the cumulative hatching probabilities and accumulated GDD values depends on how the accumulated GDD is calculated; that is, it depends on the GDD (t) profile, namely on the cardinal temperatures, Eq. (1), and the starting date of accumulation. Thus, we first calibrated our model to obtain the GDD temperature profile and a starting date of GDD accumulation that provided the best-predicted hatching probabilities for our dataset.

We selected three plausible starting dates for GDD accumulation as described above: 1^st^ of November, 1^st^ of December and 1^st^ of January. Then we calculated the optimal GDD temperature profiles, by finding the values of *T_base_, T_opt_* and *T_max-_* that gave the best fit with the observed hatching probabilities in the four field experimental sites. More precisely, we tested all the combinations with *T_base_* from 4°C to 12°C, *T_opt_* from 20°C to 28°C and *T_max_* from 30 to 42°C in steps of 0.2°C. Then, we predicted the hatching probabilities using Eq. (2) the accumulated GDDs computed with each profile and each starting date. Finally, we selected the optimal profile (the optimal values of *T_base_, T_opt_* and *T_max_*) to define the profile that minimized the error between Eq. (2) and the experimental dataset for each starting date.

### Model Validation

For each diapause breakage date considered (1^st^ of November, 1^st^ of December and 1^st^ of January), an optimal GDD temperature profile can be calculated from our dataset (the 4 experimental sites). However, we were interested in extrapolating those results from our model to other locations in Spain for which the egg-hatching dates were recorded in a six-year survey. Therefore, we conducted a validation process to find out which diapause breakage date fitted best to our six-year dataset. Thus, independent data of field observations on the occurrence of newborn nymphs in different regions of Spain during 2016-2021 were used to validate the model (Supp. Table 4). These data were obtained from systematic field surveys conducted by different researchers from public institutions in Spain (Andalucía, La Rioja, Madrid, Murcia and, Valencia) (see acknowledgments).

### Control timing of *P. spumarius* nymphs

After model validation, we used our calibrated and validated model to investigate the best timing to adopt control actions against *P. spumarius* nymphs based on field observations. To do so, we calculated the daily hatching probabilities of *P. spumarius* eggs in the Iberian Peninsula based on ERA5-Land temperature data (Muñoz Sabater 2019), simulated applications of control actions at several probability levels, and calculated the efficacy of applying a control action at a given time. Thus, the following algorithm was developed to determine the timing of control actions based on our model predictions. First, we selected a plausible range of diapause breakage dates as the starting dates for our simulations. Although we calibrated and validated our model for a particular diapause breakage date, we investigated if different dates with the same temperature profile were able to provide better results; then, we selected the ending date of each simulation as a plausible date on which no more egg hatching was expected. Afterward, we selected several probability levels to apply control actions at a given time (i.e., when any of these selected levels is crossed in any location, we simulated the application of a control action). For each diapause breakage date in each simulation, the daily hatching probabilities were computed until the end of the simulation. Then, we saved the dates at which every selected probability level (control action dates) at each location matched the available data for the presence of newborn nymphs in the field. (Supp. Table S4). Finally, the lags between the control action dates and observation dates were computed.

Each hatching probability (which matches an action date) has a given control action efficacy. It is known that nymphal development lasts for approximately 5–6 weeks until they become adults, but the developmental rates decrease when temperatures are low and could extend to at least 100 days (Weaver and King 1954, Yurtsever 2000, Bodino et al. 2019). Thus, we followed a conservative approach to compute the control action efficacies and defined successful nymph control only if the action was applied after the newborn nymphs were observed and there was a lag of less than 30 days.

According to our results, there is high variability in the hatching dates (see the results section). Thus, to overcome this intrinsic variability in the timing of egg hatching of *P. spumarius*, we developed a strategy based on deploying control actions at two different times to target the maximum number of nymphs as completely as possible. A list of combinations of two probability levels was selected (e.g., the first at 50% and the second at 90%) to apply control actions on two different dates. Moreover, the efficacies of applying control actions at a single time or at two different times were compared.

In addition, we developed an R package script to calculate the daily cumulative hatching probabilities of *P. spumarius* eggs based on suitable temperature data. This package also calculates the optimum time to apply the first and second control actions, depending on the hatching probability achieved. This package and all relevant information regarding its use can be found on the GitHub repository: https://github.com/agimenezromero/PSEggHatching.

### Data analyses

After monitoring egg-hatching in each of our four experimental sites, we explored whether the oviposition date could or could not affect the egg-hatching date within each site. In addition, after model validation, we studied the impact of relative humidity (RH) on the GDD needed for hatching. We computed RH as mean values experienced by each egg during the experimental period. To analyze the correlations between the different factors mentioned above we performed Spearman’s rank correlation tests (Benesty et al. 2009). Finally, we calculated the Wasserstein distance to measure the distance between probability distributions of GDD accumulation between each pair of field sites (two by two) (Panaretos and Zemel 2019). All statistical analyses were performed with R (R Core Team: 2021. V 4.1.3). The modeling was performed in Python programming language (Van Rossum and Drake 1994).

## RESULTS

### Nymphal emergence

A total of 435 *P. spumarius* nymphs emerged in the field assays at the four experimental sites. Newborn nymphs were detected feeding on *S. oleraceus* plants from 5-III-2021 to 1-V-2021. The first detection of newborn nymphs was recorded on 5-III-2021 at Alcalá de Henares (588 m), on 18-III-2021 at Bustarviejo (1,222 m), on 23-III-2021 at Pedrezuela (880 m) and 31-III-2021 at Mataelpino (1,086 m). Nymph emergences occurred at different moments and extended for approximately two months at all of the studied sites (Figure 2). The results for the nymphal emergence dates per field location show that the intrinsic randomness of the hatching process is quite broad, which exhibits variations of up to one month for eggs subjected to the same environmental conditions. No correlation between the oviposition date and hatching date was observed after pooling data from all of the field sites (ρ=-0.036). Furthermore, there was no or poor correlation between the oviposition date and hatching date within specific field sites: Alcalá Henares (ρ=-0.192), Bustarviejo (ρ=-0.289), Mataelpino (ρ=-0.016) and Pedrezuela (ρ=-0.316).

**Figure 2:**
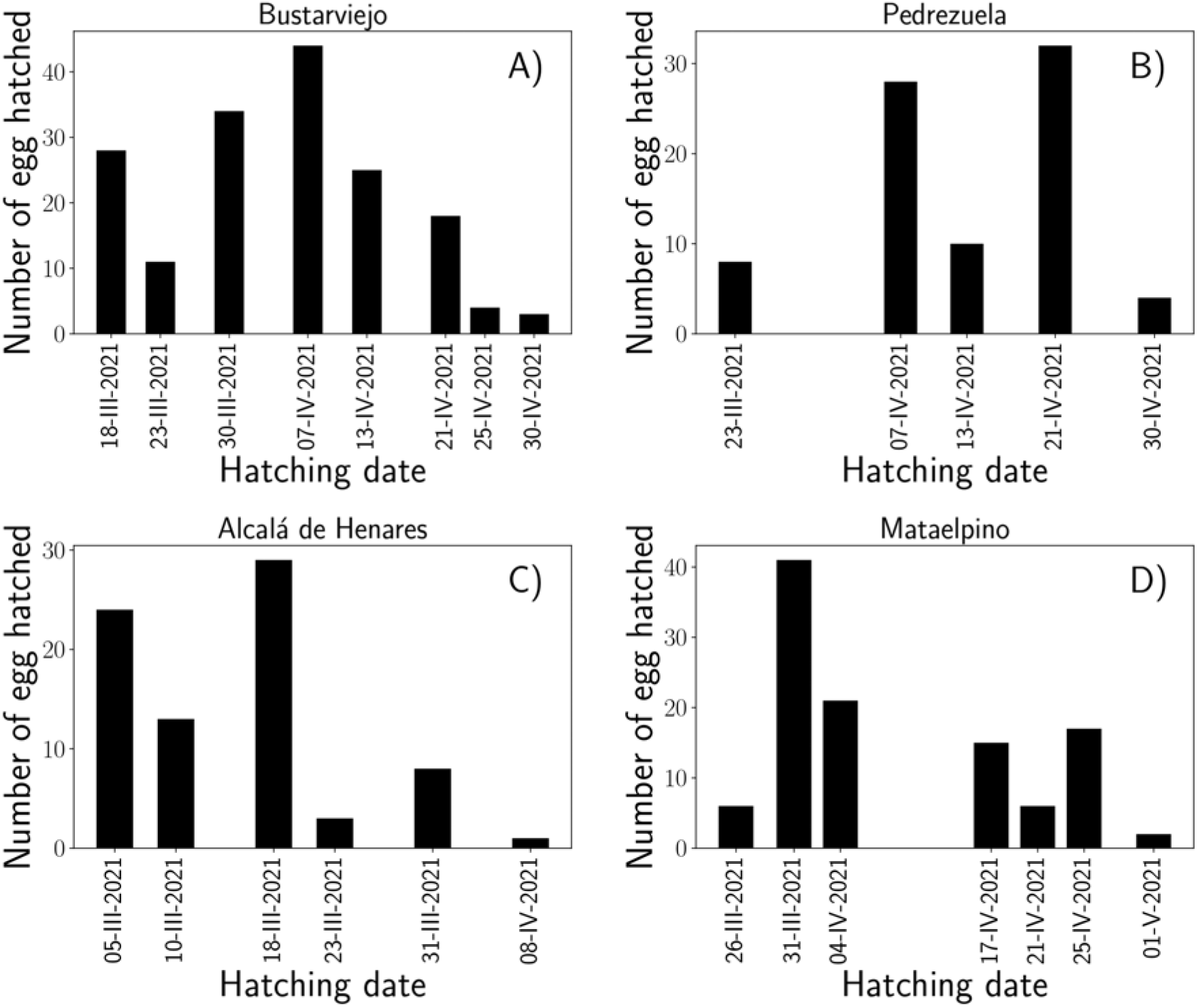
Number of eggs hatched per date at each field site. A) Bustarviejo. B) Pedrezuela. C) Alcalá Henares. D) Mataelpino.

### Model Calibration

As explained in Methods, we calibrated the GDD temperature profile assuming different diapause breakage dates: 1^st^ of November, 1^st^ of December and 1^st^ of January. We obtained an unrealistic temperature profile with high T_opt_ and T_max_ values when starting the accumulation of GDD on the 1^st^ of January (*T_base_* = 9.2°C, *T_opt_* = 27.6°C, and *Tmax* = 41.8°C) and 1^st^ of November (*T_base_* = 9°C, *T_opt_* = 27.4°C, and *T_max_* = 41.2°C). In contrast, more appropriate and realistic results were obtained when selecting the 1^st^ of December as the diapause breakage date (*T_base_* = 9.2°C, *T_opt_* = 23.4°C, and *Tmax* = 34.2°C).

As a visual example, Figure 3 shows the final result of the model after the calibration procedure for the diapause breakage date for 1^st^ December (model calibration for diapause breakage date for 1^st^ of January and 1^st^ of November are shown in Supp. Document S5). The dots represent the cumulative hatching probabilities that were retrieved from the dataset as a function of the accumulated GDD values. This accumulated GDD value was obtained using the optimal GDD profile that is shown in the inset (*T_base_* = 9.2°C, *T_opt_* = 23.4°C, and *T_max_* = 34.2°C). The black line is the best fit of Eq. (1) to the experimental data (dots) that was obtained with *k* = 4.34 and *λ* =164.86, which had a relative error of only 1%.

**Figure 3:**
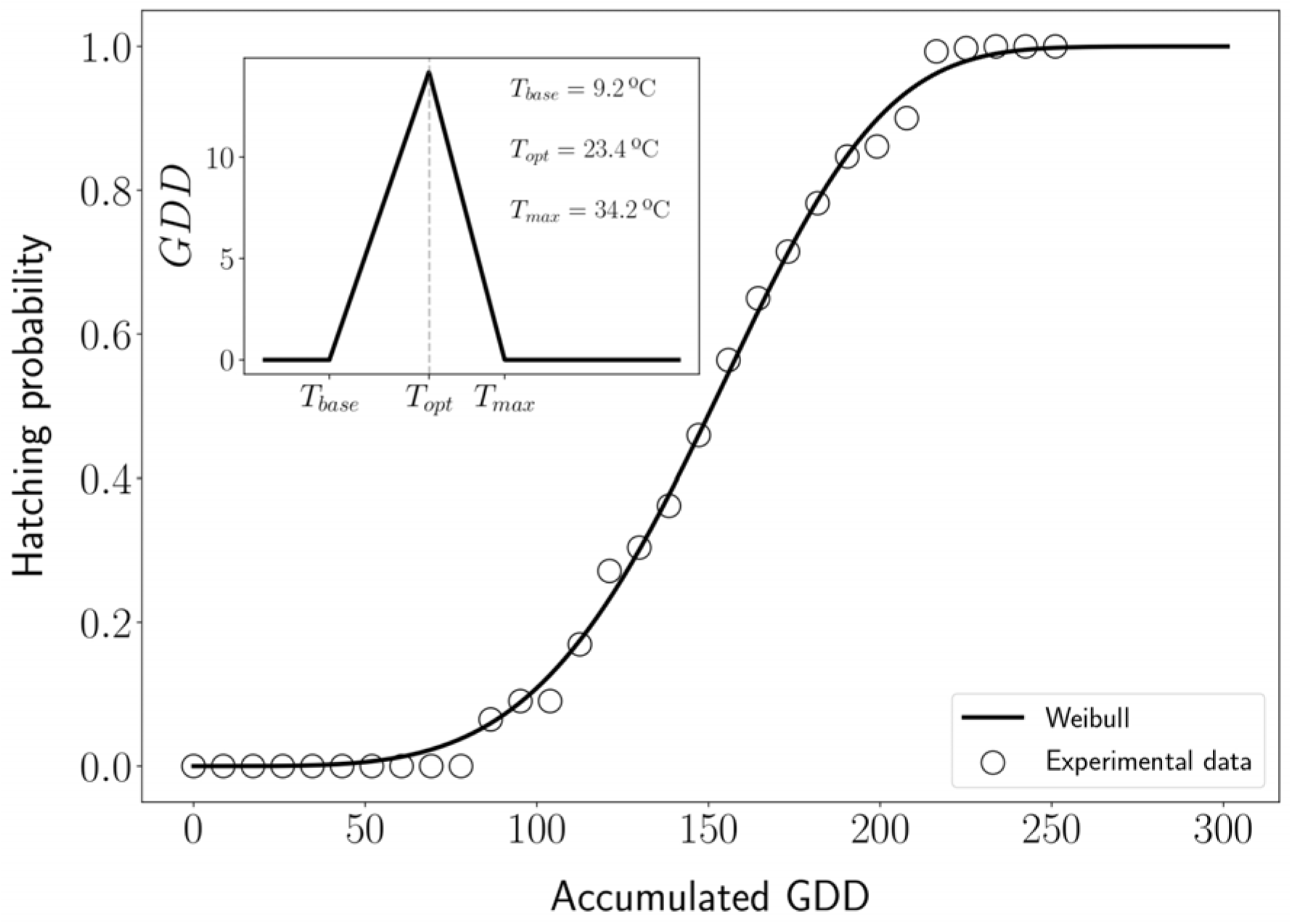
Model calibration considering diapause breakage on December 1^st^. The main figure shows the hatching probability as a function of the accumulated GDD value. Dots represent the experimental data, and the black solid line shows the best fit using Eq. (1). The inset shows the GDD profile that is used to calculate the accumulated GDD values, which yielded the best fit between Eq. (1) and experimental data.

### Model validation and predictions of egg hatching dates in the Iberian Peninsula

To validate our model, we predicted the egg-hatching probabilities in the Iberian Peninsula by using the existing dataset from field observations recorded from 2016 to 2021 in Spain (Supp. Table S4) using the calibrated model considering the three different dates of diapause breakage date. The hourly temperature data for these years were retrieved from the ERA5-Land dataset, which has a spatial resolution of 0.1° x 0.1°. Then, we computed the hatching probabilities with daily resolution and compared the predictions with the observational data from the field that were obtained from systematic surveys of newborn nymphs in different regions of Spain (Supp. Table S4). These data consisted of several records of GPS coordinates and dates where newborn nymphs were found in Spain, so we expected that our model would predict high probabilities of egg hatching (> 50%) for each date and location where newborn nymphs were observed. It was found that our model predictions were only consistent with the field data when the diapause breakage date was set to the 1^st^ of December (Supp. Video S6). When assuming the 1^st^ of January, the model predicts egg hatching too late so that nymphs are observed about a month before our model predicts high hatching probabilities (Supp. Video S7). On the other hand, when assuming November 1^st,^ the model predicts a high probability of egg hatching too early, about 2 months before field observations (Supp. Video S8). Nevertheless, considering the diapause breakage as the 1^st^ of December, some differences were also observed between model predictions and field observations from systematic surveys (Supp. Table S4) in southern Spain (latitudes below 40°) and northern Spain (latitudes above 40°). As shown in Supp. Video S6, in the South, 1^st^ instar nymphs were detected in the field, when the model predicted high hatching probabilities (above 80%), while in the North 1^st^ instar nymphs are detected when the model predicted low hatching probabilities (20%-30%).

### Impact of RH on GDD for hatching and GDD accumulation in the field sites

After model calibration and validation, considering diapause breakage on December 1^st^ we explored the impact of RH on GDD needed for hatching. As shown in Figure 4A, there is no or weak correlation between RH and GDD within field sites (Alcalá de Henares ρ= 1.0; Bustarviejo ρ= 1; Pedrezuela ρ= 0.249 and Mataelpino ρ= 0.392) and the GDD needed for hatching were similar in all field sites, except for Bustarviejo, which deviates from the general pattern. We found that the GDD needed for hatching was independent of field sites. As shown in Figure 4B, the probability density function of egg hatching as a function of accumulated GDD is very similar in all field sites, except Bustarviejo. The boxplot of Figure 4C again shows that the GDD needed for hatching is similar in all locations except Bustarviejo. More quantitatively, Figure 4D shows the Wasserstein distance among the distributions, showing that Bustarviejo is the only one different from the others. Finally, temperatures and RH registered in the four field locations followed a similar pattern. Temperatures were lower in winter and higher during autumn and spring, while RH was higher during autumn and winter, being lower in March and starting to increase in April (Supp. document S9).

**Figure 4:**
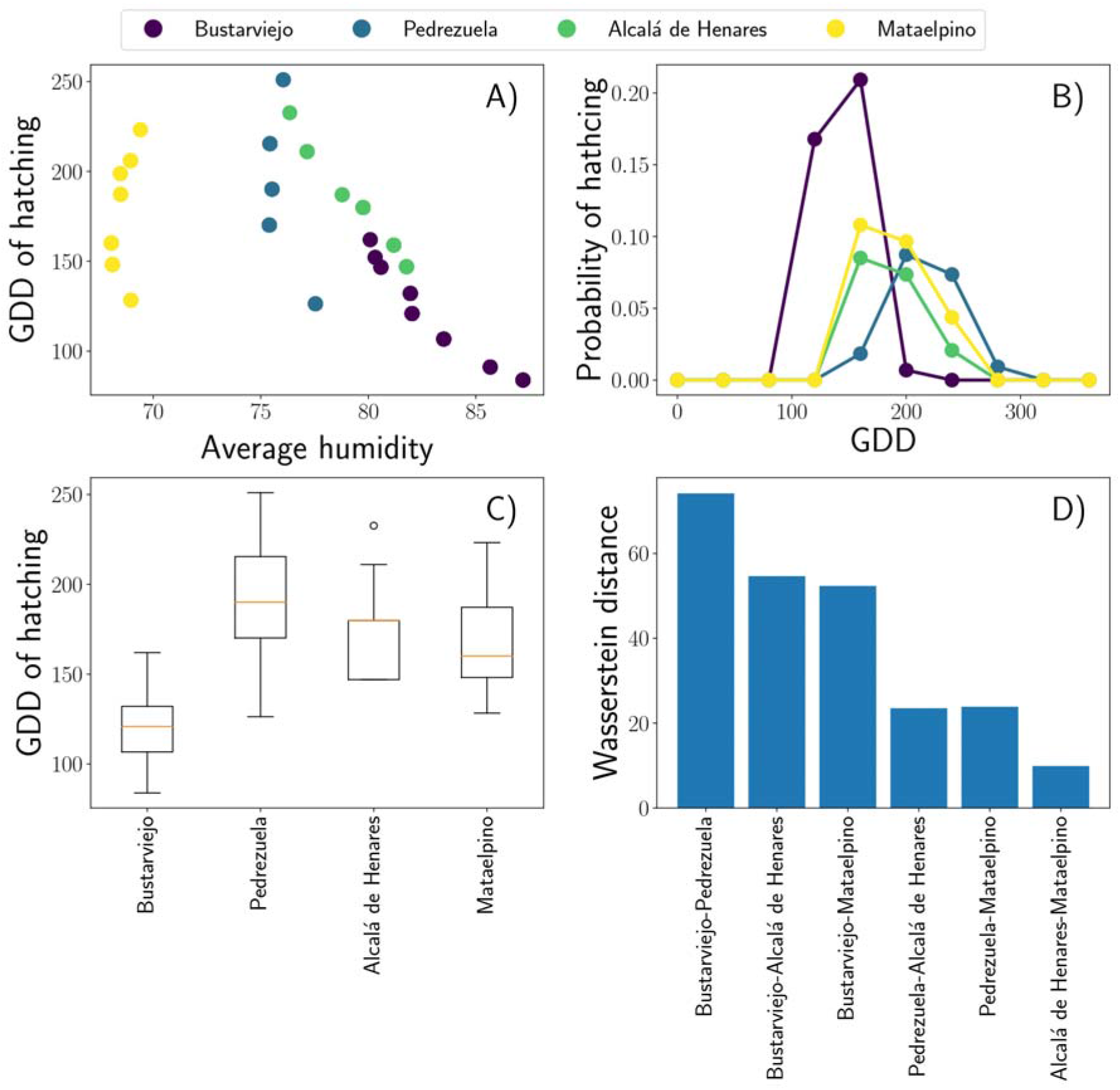
A) GDD accumulation metric at the moment of egg hatching versus the RH values at the experimental locations. Each data point corresponds to one egg, and the colors correspond to different locations. B) Probability density function of egg hatching as a function of accumulated GDD C) Distribution of GDD needed for egg hatching on each field site. The horizontal orange lines represent median values. Boxes extend from the 25th to the 75th percentile of each group’s distribution of values, and vertical extending lines indicate the range of values. D) Wasserstein Distance (WD) between probability distributions of GDD accumulation between each pair of field sites (two by two).

### Decision-support tool to determine the best timing for controlling *P. spumarius* nymphs

After the model validation, we calculated the precise timing for controlling *P. spumarius* nymphs most efficiently in different regions in Spain. The following initial conditions were fixed according to our results: 1) We selected a plausible range of diapause breakage dates: we selected the range 1^st^ of November to 1^st^ of January 2). The end date of each simulation was based on the field observations from the systematic surveys (2016-2021) (Supp. Table S4). Because the latest newborn nymphs were found on 27-V-2021 we assumed that the end date of each simulation was June 1^st^. With this algorithm in mind, the optimal dates for applying control actions (control timing) were selected to maximize the defined efficacy, which is the maximum percentage of targeted nymphs. In addition, we considered all possible date combinations for taking certain control actions and compared these simulations with all of the hatching observations from both our assays and the systematic surveys (Supp. Table S4). This maximum efficacy changed depending on the diapause breakage date (Figure 5 A-B). Consistent with the previous validation, it was found that the model accuracy is maximized if diapause breakage is considered to occur in December. Interestingly, we found that assuming diapause breakage in mid-December instead of 1^st^ December is a more robust choice. In this way, if the diapause breakage date slightly varies for a given particular year, we expect to maintain high efficacy (Figure 5 A-B). Furthermore, to optimize the control actions and to target the maximum number of nymphs as effectively as possible, a control strategy (control timing) applied at a single date was compared to a two-date application strategy. Our results clearly show that the best strategy consists of applying control actions at two different times to target the maximum nymphal population (Figure 5 C-D). In contrast, the maximum efficacy achieved by applying only one control action is below 50% in both northern and southern Spain. According to our results, for northern Spain (latitudes above 40°N) the first control actions should be taken when the accumulated egg-hatching probabilities reach 40% and the second when they reach 90%. On the other hand, for southern Spain (latitudes below 40°N), the first control actions should be taken when the accumulated egg-hatching probability reaches 30% and the second when it reaches 90% (Figure 5).

**Figure 5:**
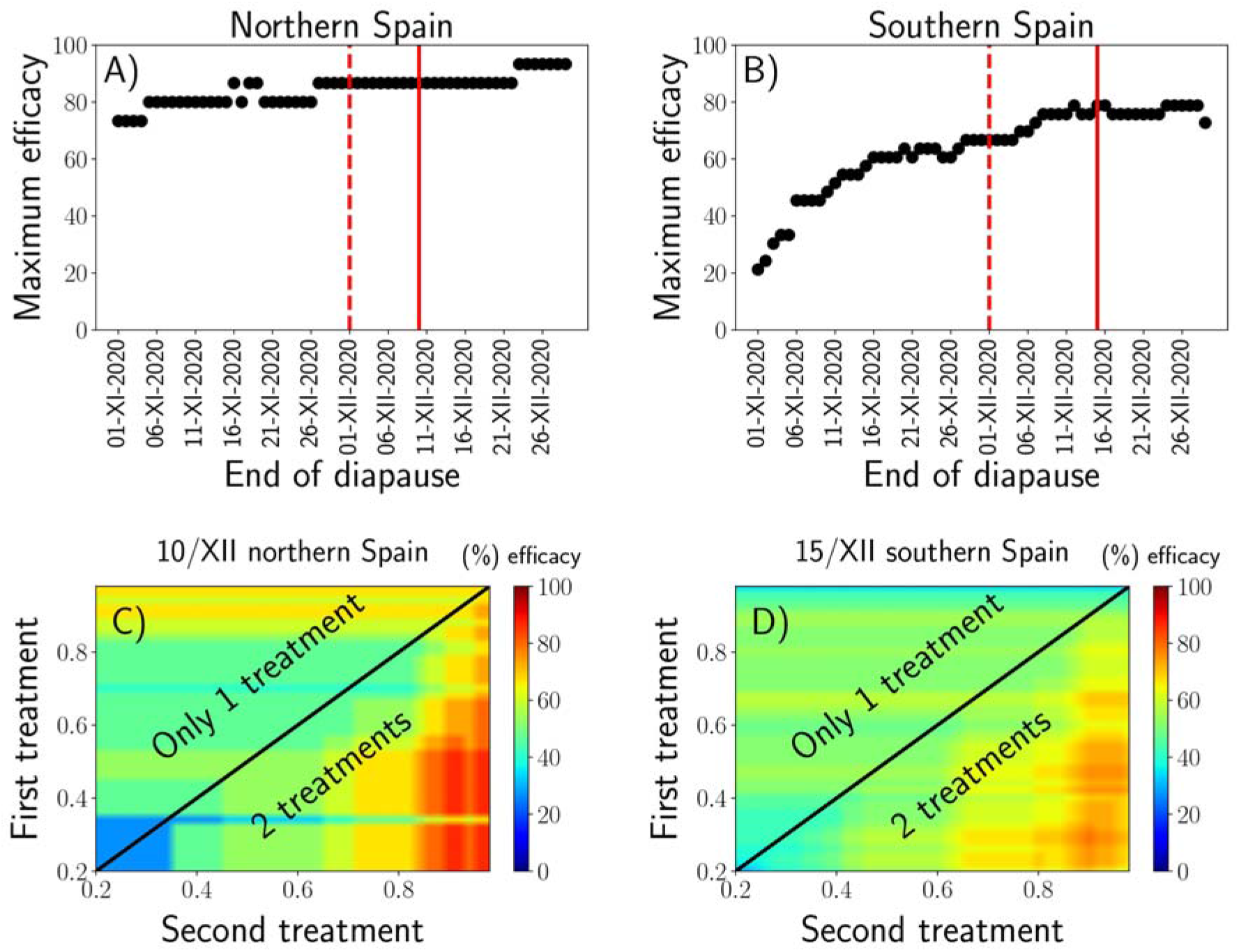
(A and B) Maximum control action efficacies for each diapause breakage date from the selected range of dates to develop the algorithm (from November 1^st^ to December 30^th^). The dashed line shows the diapause breakage selected for model development (December 1^st^) and the solid line indicates the diapause breakage date to achieve the maximum efficacy (December 10^th^ in the north and December 12^th^ in the south). A) Efficacy in northern Spain (latitude >40°N). B) Efficacies in southern Spain (latitude < 40°N). (C and D) Examples of the efficacies of applying a control action depending on the egg hatching probability after applying one or two treatments (C) in northern Spain when considering diapause breakage on December 10^th^ and (D) in southern Spain when considering diapause breakage on December 15^th^. This analysis was repeated for all diapause breakage dates to obtain the maximum efficacies shown in (A and B).

## DISCUSSION

Growing degree-day (GDD) models, or degree-day models (Herms 2004), are often used in agronomy to determine the correlations between development and temperature to forecast the times when biological events will occur. In addition, these models can be used as decision-making tools to establish the best times to target specific pests (control timing) and optimize control measures, such as cultural (e.g., tillage) or chemical (e.g., insecticides) control measures to increase their effectiveness and to mitigate their side effects (Herms 2004).

Some models have aimed to forecast the development of *P. spumarius*. In the study performed by Bodino et al. (2019), the authors estimated the base temperatures from assays performed at fixed temperatures; the model was constructed based solely on a *T_base_* threshold. On the other hand, Chmiel and Wilson calculated the *T_base_* and *T_max_* thresholds for development based on the lowest coefficient of variation (Arnold 1959) and fitted a linear regression function. In addition, other studies on different insect species based on linear GDD functions that used only the *T_base_* threshold have been proposed (Johnson et al. 1998; Bodino et al. 2019). Conversely, we obtained the temperature thresholds directly from experimental data obtained from field experiments under natural conditions and constructed a model based on *T_base_, Tmax* and *T_opt_*. Our approach allows us to indirectly account for the effect of temperature fluctuations on the developmental rate, which is automatically incorporated into the model within the best-fit parameters. The temperature has significant nonlinear effects on insect development, which is slow when the temperatures approach either the upper or lower developmental thresholds (Murray 2020). For this reason, other studies based on nonlinear approximations have also been developed (Logan et al. 1976, Lactin et al. 1995, Briere et al. 1999). In our work, we provided a mathematical approach based on temperature effects on the metabolic rates of ectotherms (Gillooly et al. 2001) to model the nonlinear development rates with a multilinear function. The model calibration and validation further support our approach of using a multilinear function to calculate the GDD metric and validate the direct use of field data to accurately determine the temperature thresholds. Similar to Bodino et al. (2019), we installed data loggers inside the cages where the eggs were placed to obtain similar temperature and humidity data as those experienced by the eggs, while Chmiel and Wilson (1979) obtained the temperature data from meteorological stations. In the review performed by Bonhomme (2000), it was stated that temperatures should be measured at the precise sites where studies are conducted instead of using temperatures obtained from meteorological stations, since microclimate variations could be important, and the actual temperatures could differ from those obtained at meteorological stations.

One of the critical components of using GDD models to predict insect phenology is the determination of the starting points for degree-day accumulations (Kim et al. 2020). Ideally, it should be set up when insect development begins. Egg development of the meadow spittlebug starts after a winter diapause. In the GDD model introduced by (Chmiel and Wilson 1979), they arbitrarily set the GDD accumulation starting point at the 1^st^ of January. However, the precise hours or days needed under a given temperature -cold requirement- and the precise length of the winter diapause of *P. spumarius* is unknown. Initially, we assumed for our model that the 1^st^ of January was the date when the diapause ended and when GDD accumulation started. However, when such an assumption was made, the temperature profile to compute GDD’s (T_min=9.2, T_opt=27.6, T_max=41.8°C) appeared to be unrealistic because the maximum temperature yielded by the model (T_max=41.8°C) is well above the biological limit for survivorship of the insect, and the optimal is also too high (T_opt=27.6). Secondly, when validating the model with observations of 1^st^ instar nymph emergence in several locations in Spain during 6 consecutive years, the model predictions did not match field observations (Supp. Video S7). Thus, we assumed that diapause breakage should occur earlier than January 1^st^ in our region and tested December 1^st^ and November 1^st^ as the starting dates for GDD accumulation. When we tested November 1^st^ as the diapause end date, the model also yielded an unrealistic optimal and maximum temperature profile (*T_base_* = 9°C, *T_opt_* = 27.4°C, and *Tmax* = 41.2°C) and also matched rather bad the results of 1^st^ instar nymphal emergence in Spain. When we tested December 1^st^ as the starting date for GDD accumulation the temperature profile (T_min=9.2°C, T_opt=23.4, T_max=34.2) was more realistic and the model predictions matched the predictions of 1^st^ instar nymph emergence in Spain and were much more convincing (Supp. Video S6).

The main reason for the discrepancy between the dates when the diapause ends in the study conducted by Chmiel and Wilson in 1979 and the earlier date that we assumed in our study is presumably related to the striking differences in the winter climate between the two regions. While the former study was conducted in West Lafayette, Indiana, USA where winter temperatures are extremely low, our study was conducted in the Iberian Peninsula where temperatures are milder thus, we presume that the cold requirement for diapause breakage of *P. spumarius* would also differ between the two regions. The induction and termination of winter diapause are crucial components of the life cycle of insect species subjected to diapause (Posledovich et al. 2015, Tougeron 2019) and a geographic variation in diapause duration depending on temperature has been previously reported by several authors (e.g. Schmidt et al. 2005, Salman et al. 2019). It is also known that autumn conditions may be a major driver of local adaptation of diapause traits (Masaki 2002). Insect populations often differ in the duration of diapause, which is likely to vary depending on the climate conditions, leading to local adaptation in the insect life cycle (Lindestad et al. 2020). For example, in the study conducted by Chuche and Thiéry (2009), they reported that the hatching dynamics of the leafhopper *Scaphoideus titanus* were cued by cold temperatures experienced during winter and may explain the variations of their population dynamics depending on the latitude. Therefore, we assumed that *P. spumarius* populations that occur in Spain are adapted to milder winters and thus, they require fewer hours of cold for diapause breakage. Furthermore, the differences observed between our model predictions and field observations of egg hatching in northern (latitude >40°) and southern (latitude<40°) Spain (Supp. Video S6), could be also explained by differences in cold temperatures experienced throughout winter (Witsack 1973) or different spittlebug populations may have different temperature profiles as an adaptation to regional climate. Our results suggest that the diapause requirements are different between spittlebug populations depending on the regional climate conditions. However, further studies should be performed for determining the precise cold requirement (hours or days under a given temperature) for winter diapause for different spittlebug populations.

In the present study, we also explored the impact of humidity on the egg-hatching of *P. spumarius*. According to our results, no correlation was found between GDD needed for hatching and RH in the field sites (Figure 4A). Nevertheless, several authors suggest that humidity has a strong influence on the life cycle of spittlebugs (Weaver and King 1954, Whittaker 1970, Ossiannilsson 1981). Therefore, further studies are needed to understand the role of humidity on the egg-hatching of *P. spumarius*.

According to our results, the emergence of *P. spumarius* neonate nymphs extended for about two months (Figure 2) and the GDD accumulation was similar in all the field sites except Bustarviejo (Figures 4B, C and D). The dates and the duration of egg hatching in the Iberian Peninsula in the present study are similar to the observations of several authors in Mediterranean crops (Morente, Cornara, Plaza, et al. 2018, Bodino et al. 2019). It is worth noting that there exists a bias between the detection of 1^st^ instar nymphs during our field experiments and the real date of egg hatching since the egg masses were revised only once a week. Therefore, the slight differences observed in data distribution of GDD accumulation between Bustarviejo and the rest of the field sites may be caused by this bias. Due to this bias, we measured the distance between probability distributions of GDDs between field sites with the Wasserstein distance (Panaretos and Zemel 2019) instead of comparing means or median with statistical tests (e.g. ANOVA or Kruskal Wallis), to avoid wrong interpretation of the data. Nevertheless, the temperature profile considering diapause breakage on December 1^st^ (T_min=9.2°C, T_opt=23.4, T_max=34.2) is convincing and the model predictions are consistent with the field observations (Supp. Table 4). Slight differences in *P. spumarius* phenology between field sites have been also previously reported in Italy (Bodino et al. 2019).

Applications of predictive analysis in ecology seek to establish models of certain problems that need to be solved. Indeed, before applying any control actions, the best time of application, control timing, should be defined. The timing for applying pest control actions should be established by developing decision-making systems that are defined by risk thresholds (Pedigo 1986, Lima et al. 2019, Dean et al. 2021).

One of the pivotal factors in the epidemiology of *X. fastidiosa* diseases is the vector’s ability to reach high population densities (Purcell 1975, Purcell et al. 1979, Almeida and Purcell 2003), which could counterbalance their low transmission efficacy (Daugherty and Almeida 2019). Moreover, it is essential to apply control actions at the appropriate developmental stage. Since *P. spumarius* nymphs have limited mobility and only adults contribute to the spread of *X. fastidiosa* to woody hosts, control actions should be focused on the preimago (nymphal) phase (Cornara et al. 2016, 2018). Therefore, control-timing actions should be conducted when nymphal populations reach their maximum densities. The emergence of *P. spumarius* nymphs is inherently a stochastic process; thus, it does not occur on a fixed date (Morente, Cornara, Plaza, et al. 2018, Bodino et al. 2019). In addition, the nymphal stage passes through five instars, and their development takes 5–6 weeks until adults emerge (Weaver and King 1954, Yurtsever 2000, Bodino et al. 2019). Given the time window for nymphal emergence and development, our predictions suggest that controlling nymphs at two different dates would target the highest percentages of nymphal populations present in the field. Moreover, the first control actions in the north should be taken when the accumulated egg-hatching probability reaches 40% and the second when it reaches 90%. For southern Spain, the first control actions should be taken when the accumulated egg-hatching probability reaches 30% and the second when it reaches 90%.

In the present study, we provided an R package script for practical use, to compute GDDs in a given location based on the local temperatures, and in turn, the model can determine the hatching probability at a given site and date. Given an input dataset of hourly temperatures in a precise location from the starting date of GDD accumulation to a given date where the model is run, the package provides the current probability level for egg hatching at that given date and precise location. So, further explained, first, the user should somehow obtain hourly temperature data starting on the 1^st^ of December. Then, each day after the 1^st^ of December, the user could introduce this CSV (comma-separated values) data file containing up-to-date hourly temperature into the R program and the output would be the probability level that has been achieved up to that given date. If for that current date the probability reaches the first or second threshold the program calculates the best timing to apply the treatment.

After the detection of *X. fastidiosa* in Europe, great efforts have been devoted to developing containment strategies. The core of such strategies is the control of *P. spumarius* through the management of ground cover, which aims at targeting resident nymphal populations by mechanical means (e.g., tillage) or by the application of pesticides against vectors (EU 2020/1201, 2020). Nevertheless, both practices, when continuously repeated, can provoke serious side effects that pose risks to biodiversity, ecosystem services and human health; indeed, frequent and drastic tillage in dry Mediterranean ecosystems may lead to negative impacts on soil fertility, soil erosion and desertification (Biondi et al. 2012, Kairis et al. 2013). Therefore, degree-day-based models could be implemented under IPM programs to apply control actions promptly. This will decrease the amount of undesirable side effects to the environment and optimize the control of insect pests, including vectors of plant diseases such as *P. spumarius*. The developed model could be used as a decision-making tool to make recommendations regarding the best timing to adopt certain control actions against the main vector of *X. fastidiosa* and to reduce the risk of disease spread. Furthermore, these fitted models can provide estimates of *P. spumarius* biology, and potentially give valuable knowledge in future studies to predict biological aspects of the life cycle of other insect pests.

## ACKNOWLEDGMENTS

The authors would like to acknowledge our colleagues María Plaza and Marcos Ramírez for their help in the field experiments and maintaining the insect colonies. In addition, we acknowledge Martin Godefroid for his contribution to the initial experimental design. We are also thankful to Antonio Montserrat (IMIDA, Murcia, Spain), José Luis Ramos Sáez de Ojer (Consjería Agricultura y Ganadería, La Rioja, Spain), Jose Manuel Durán and Manuel Ruiz Torres (Junta de Andalucía, Spain) for their help with the systematic survey observations used for model validation. The work was funded by the Ministerio de Ciencia e Innovación under grant AGL2017-89604-R and fellowship PRE2018-083307 (CL, MM, AM and AF) and Grants RTI2018-095441-B-C22 (SuMaEco) and PID2021-123723OB-C22 (CYCLE) funded by MCIN/AEI/10.13039/501100011033 and by “ERDF A way of making Europe” (AGR and MAM) and Grant MDM-2017-0711 (María de Maeztu Excellence Unit) funded by MCIN/AEI/10.13039/501100011033 (AGR and MAM).

## SUPPORTING MATERIAL LEGENDS

**Figure S1.** A) Petri dishes with holes for drainage where the pine needles with egg masses were placed. B) Data logger placed inside one of the cages used during the field assay.

**Figure S2:** Scheme of the assay. A) Cylindrical mesh cages where the eggs were maintained at different field points. B) Cellulose acetate sheets that separate each cage into four and *S. oleraceus* plants planted in the ground. C) Egg mass distributions inside the cages at each field point. Each circle represents one cage. Each portion of the circle corresponds to one division of the cage.

**Document S3.** Explanation of the temperature response function as a generalization of the linear approximation approach.

**Table S4.** Systematic surveys conducted in Spain from 2016-2021.

**Document S5:** Model calibration considering diapause breakage on January 1^st^ and November 1^st^. Figures show the hatching probability as a function of the accumulated GDD value. Dots represent the experimental data, and the black solid line shows the best fit using Eq. (1). The inset shows the GDD profile that is used to calculate the accumulated GDD values, which yielded the best fit between Eq. (1) and experimental data.

**Videos S6, S7 and S8.** Example of the evolution of the hatching probability in the Iberian Peninsula from 2017 to 2018 when using temperature data from ERA5-Land. White dots correspond to nymphs that were observed in those given locations (coordinates) on a given date. S6) Evolution of the hatching probability considering diapause breakage on the 1^st^ of December. S7) Evolution of the hatching probability considering diapause breakage on the 1^st^ of January. S8) Evolution of the hatching probability considering diapause breakage on the 1^st^ of November.

**Document S9.** Monthly mean (± SD) temperatures (°C) and RH (%) measured in the four field sites in Bustarviejo, Mataelpino, Pedrezuela and Alcalá Henares.

## Notes

https://github.com/agimenezromero/PSEggHatching

### Competing Interest Statement

The authors have declared no competing interest.

