## Supplementary figures and images for "Degree-day-based model to predict egg hatching of *Philaenus spumarius* (Hemiptera: Aphrophoridae), the main vector of *Xylella fastidiosa* in Europe"

### Supporting Figure S1

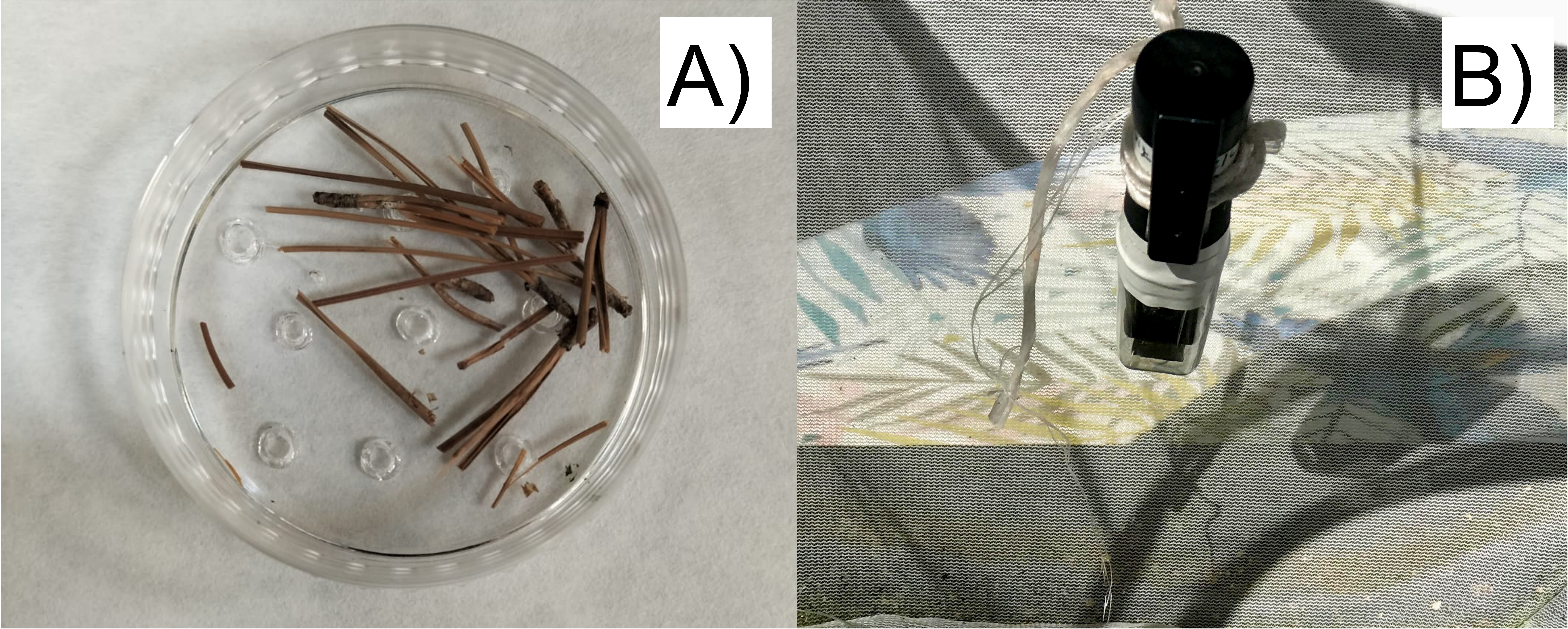

### Supporting Figure S2

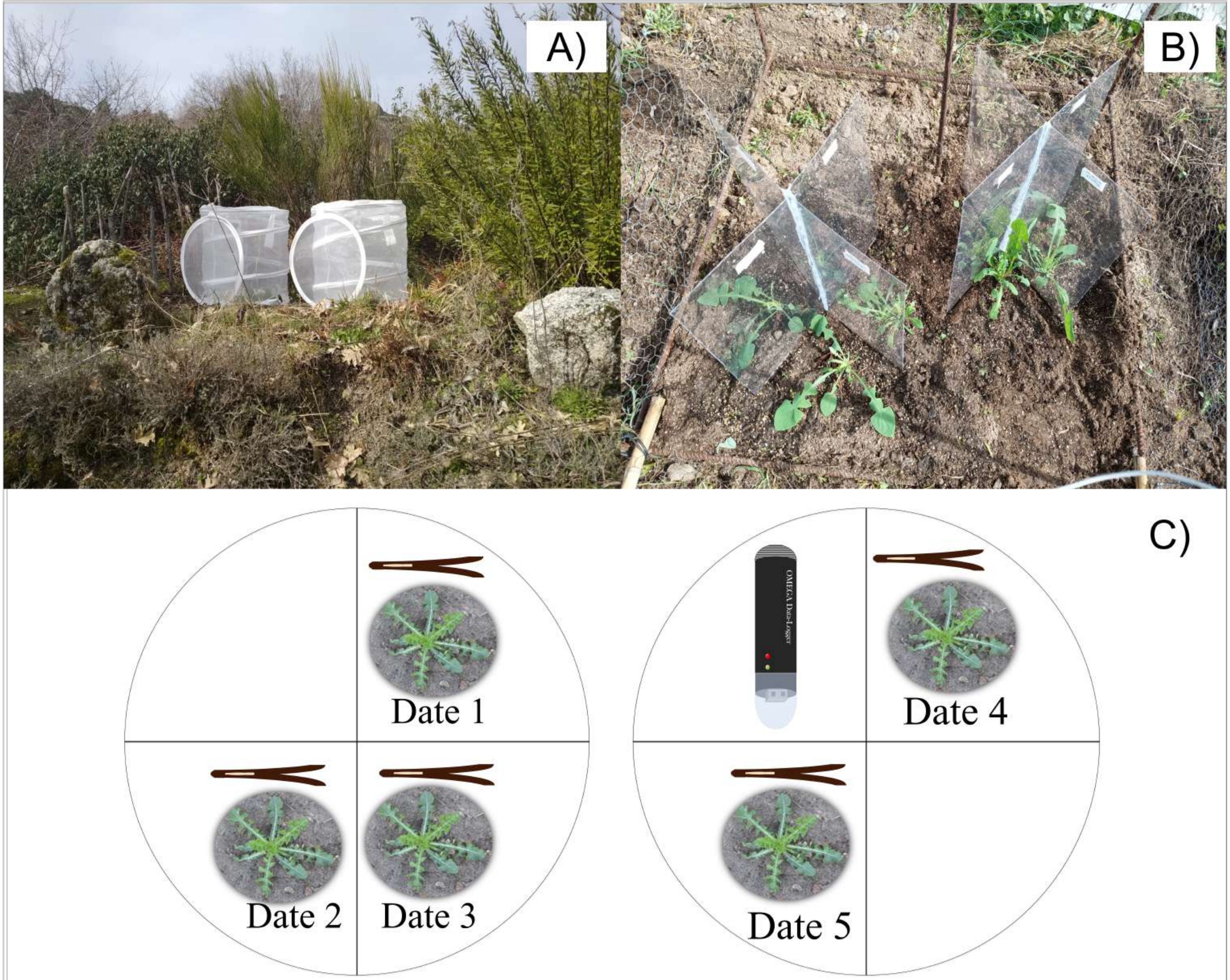
