## Supporting Document S3 for "Degree-day-based model to predict egg hatching of *Philaenus spumarius* (Hemiptera: Aphrophoridae), the main vector of *Xylella fastidiosa* in Europe"

### Supplementary Material

#### A Multi-linear approximation for $GDD$ computation

Many authors have studied the dependence of insect development on temperature, finding a non-monotonic dependence with a minimum temperature $T_{min}$, below which there is no development, and a maximum temperature $T_{max}$, above which development stops. Growth is maximum for a certain temperature, $T_{opt}$. So, a number of functional forms have been suggested to capture this unimodal behavior (see, e.g., Sharpe and DeMichele 1977; Schoolfield et al. 1981; Logan et al. 1976; Lactin et al. 1995; Briere et al. 1999). These functional forms yield smooth curves, but depend on a large number of parameters.

In this work we devise a simplified form of the temperature profile used to compute the Growing Degree-Days metric. The multi-linear generalization of the $GDD$ function is based on an approximation to an Arrhenius' Law description of temperature effects on ecthotherms (and other poikilothermal organisms). In (Gillooly et al. 2001) the authors derived a relationship, based on principles of biochemical kinetics and allometry, that characterizes the effects of temperature and body mass on metabolic rate, such that, in a suitable range, the dependence is approximately exponential on $1/T$, in the form of Arrhenius' Law.

The mathematical form of the Arrhenius' Law dependence between the growth rate $k$ and the absolute temperature $T$ reads as follows,

$$k = A\exp(-E/T) \ , \tag{1}$$

where $A$ is a pre-exponential factor and $E$ an activation energy in units of the Boltzmann constant $k_B$. The original use of this equation is for the rate constant of a chemical reaction that increases monotonically with $T$, and so $E > 0$. To account for the unimodal form of the insects development rate based on this first principles, one can consider two Arrhenius functions with opposite signs in the activation rate,

$$k = A_1\exp(-E_1/T) + A_2\exp(+E_2/T) \ , \tag{2}$$

where $E_1 > 0$ and $E_2 < 0$, as has been suggested in Begasse et al. 2015, based on the the transition state theory of Eyring 1935, which includes a term accounting for reversible protein denaturation at high temperature F. H. Johnson and Lewin 1946.

Now let us denote by $t$, the temperature in Celsius, $t = T - b$ with $b = 273.15$. Within the typical insect development temperature range (say 0-40 °C) $t$ is small respect to $b$, the absolute (Kelvin) temperature. Thus, the two exponents in Eq. (2) can be approximated as,

$$k = A\exp\left(-\frac{E}{b+t}\right) = A\exp\left(-\frac{E}{b(1+t/b)}\right) \approx A\exp\left[-\frac{E}{b}\left(1-\frac{t}{b}\right)\right] = A\exp\left(-\frac{E}{b}\right)\exp\left(\frac{E}{b^2}t\right) \approx$$
$$A\exp\left[-\frac{E}{b}\right]\left(1+\frac{E}{b^2}t\right) = A\exp\left(-\frac{E}{b}\right) + A\frac{E}{b^2}\exp\left(-\frac{E}{b}\right)t = B + Ct \ , \tag{3}$$

where we assume that $t/b = t/273.15 \ll 1$ and $(Et/(b^2) = (Et)/(273.15^2) \ll 1$, whereas $B$ and $C$ are constants. In particular, $C > 0$ if $E > 0$ fits the region before the maximum in which the growth rate increases, while $C < 0$ if $E < 0$ fits the region after the maximum where $k$ decreases. The positive/negative sign stems from the the coefficient of the linear term in $t$, $E/b^2$.

Thus, each exponential in Eq. (2) can be expressed with a simple straight line, valid in the typical development temperature range of insects (see Fig. 9). If experimental data on the insect development rate is available, this approach can be extended by adding more exponential terms in Eq. (2), to fit the experimental data with a multi-linear dependence on temperature.

With this we give a ground to use a multi-linear temperature response function to compute the $GDD$,

$$f(T) = \begin{cases} 0 & \text{if} \quad T < T_{\text{base}} \\ T - T_{\text{base}} & \text{if} \quad T_{\text{base}} \leq T < T_{\text{opt}} \\ m \cdot T + n & \text{if} \quad T_{\text{opt}} \leq T < T_{\text{max}} \\ 0 & \text{if} \quad T \geq T_{\text{max}} \end{cases} \quad \text{with } m = -\frac{T_{\text{opt}-T_{\text{base}}}}{T_{\text{max}} - T_{\text{opt}}},\ n = -m \cdot T_{\text{max}} \ , \tag{4}$$

412 to account for the unimodal dependence of insect development. Then, the accumulated $GDD$ in a certain period is given by

$$GDD = \int_{t_0}^{t_f} f(T)\,\mathrm{d}t \ . \tag{5}$$

414 The $GDD$ function Eq. (4) depends only on 3 parameters to be fitted, $T_{min}$, $T_{opt}$ and $T_{max}$.

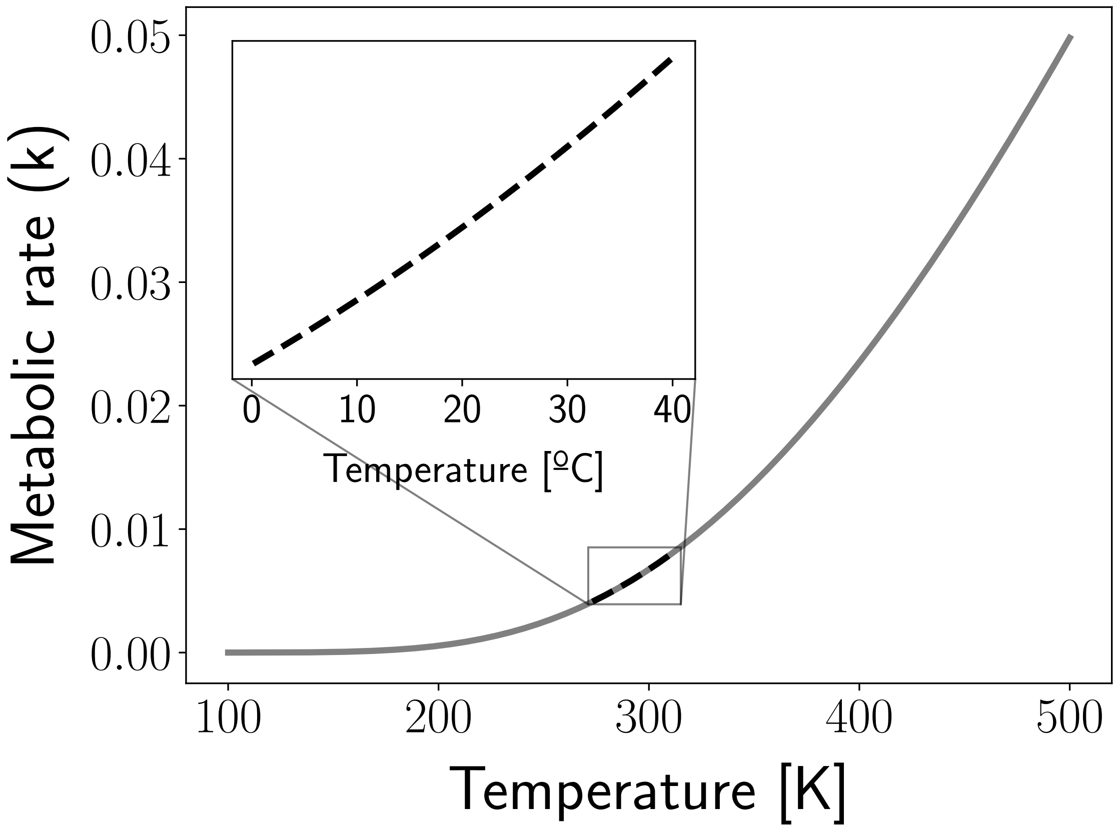


**Figure 9:** Schematic representation of the linear regime in the Arrhenius function.
