## Supporting Table S4 for "Degree-day-based model to predict egg hatching of *Philaenus spumarius* (Hemiptera: Aphrophoridae), the main vector of *Xylella fastidiosa* in Europe"

| Date of emergence | Field location | Region | Nymphal stage | Lat | Lon | Year | Observer | Origin of the data |
|---|---|---|---|---|---|---|---|---|
| 3-feb-2016 | Puerto Moral | Huelva | 1 | 37.87485635 | -6.464657663 | 2016 | Jose Manuel Durán | Systematic sampling |
| 11-feb-2016 | Castillo de las Guardas | Sevilla | 1 | 37.68167052 | -6.252431569 | 2016 | Jose Manuel Durán | Systematic sampling |
| 18-feb-2016 | Osuna | Sevilla | 2 | 37.15849409 | -5.139528214 | 2016 | Jose Manuel Durán | Systematic sampling |
| 10-mar-2016 | Constantina | Sevilla | 2 | 37.86131266 | -5.57711167 | 2016 | Jose Manuel Durán | Systematic sampling |
| 29-mar-2016 | Morón | Sevilla | 1 | 37.10039243 | -5.377321998 | 2016 | Jose Manuel Durán | Systematic sampling |
| 15-feb-2017 | Castillo de las Guardas | Sevilla | 1 | 37.6688293 | -6.251865571 | 2017 | Jose Manuel Durán | Systematic sampling |
| 7-mar-2017 | El Ronquillo | Sevilla | 1 | 37.65966196 | -6.156940406 | 2017 | Jose Manuel Durán | Systematic sampling |
| 9-mar-2017 | Algodonales | Cádiz | 2 | 36.88516931 | -5.451834299 | 2017 | Jose Manuel Durán | Systematic sampling |
| 14-mar-2017 | Constantina | Sevilla | 1 | 37.862028 | -5.57736391 | 2017 | Jose Manuel Durán | Systematic sampling |
| 14-mar-2017 | Osuna | Sevilla | 2 | 37.15982699 | -5.139081596 | 2017 | Jose Manuel Durán | Systematic sampling |
| 26-feb-2018 | Castillo de las Guardas | Sevilla | 2 | 37.66339545 | -6.247655085 | 2018 | Jose Manuel Durán | Systematic sampling |
| 13-mar-2018 | Constantina | Sevilla | 2 | 37.861944 | -5.576111 | 2018 | Jose Manuel Durán | Systematic sampling |
| 7-abr-2018 | Osuna | Sevilla | 3 | 37.1594444 | -5.1394 | 2018 | Marina Morente | Systematic sampling |
| 17-abr-2018 | Arnedo | La Rioja | 1 | 42.0012839 | -1.99117916 | 2018 | Marina Morente | Systematic sampling |
| 19-abr-2018 | Morata Tajuña | Madrid | 2 | 40.231134 | -3.450696 | 2018 | Marina Morente | Systematic sampling |
| 20-abr-2018 | Alberite | La Rioja | 2 | 42.4183333 | -2.44083 | 2018 | Marina Morente | Systematic sampling |
| 20-abr-2018 | Rodezno | La Rioja | 1 | 42.5291667 | -2.82861 | 2018 | Marina Morente | Systematic sampling |
| 27-ene-2020 | Cartagena | Murcia | 1 | 37.611866 | -0.750929 | 2020 | Antonio Montserrat | Systematic sampling |
| 11-feb-2020 | Alhama de Murcia | Murcia | 2 | 37.838723 | -1.467839 | 2020 | Antonio Montserrat | Systematic sampling |
| 11-feb-2020 | El Berro | Murcia | 1 | 37.887945 | -1.493203 | 2020 | Antonio Montserrat | Systematic sampling |
| 11-feb-2020 | Valle Perdido | Murcia | 1 | 37.927495 | -1.148376 | 2020 | Antonio Montserrat | Systematic sampling |
| 18-feb-2020 | Jumilla | Murcia | 2 | 38.442847 | -1.316243 | 2020 | Antonio Montserrat | Systematic sampling |
| 18-feb-2020 | Jumilla | Murcia | 1 | 38.492305 | -1.189841 | 2020 | Antonio Montserrat | Systematic sampling |
| 18-feb-2020 | Yecla | Murcia | 1 | 38.618762 | -1.127371 | 2020 | Antonio Montserrat | Systematic sampling |
| 16-abr-2020 | Bustarviejo | Madrid | 3 | 40.691827 | -3.767162 | 2020 | IVPP Research group | Systematic Sampling |
| 16-abr-2020 | Bustarviejo | Madrid | 2 | 40.691827 | -3.767162 | 2020 | IVPP Research group | Systematic Sampling |
| 16-abr-2020 | Colmenar Viejo | Madrid | 2 | 40.691917 | -3.767194 | 2020 | IVPP Research group | Systematic Sampling |
| 16-abr-2020 | ICA | Madrid | 1 | 40.43961581 | -3.687287572 | 2020 | IVPP Research group | Systematic Sampling |
| 17-abr-2020 | IMIDRA | Madrid | 3 | 40.521133 | -3.290865 | 2020 | IVPP Research group | Systematic Sampling |
| 20-abr-2020 | IMIDRA | Madrid | 1 | 40.521133 | -3.290865 | 2020 | IVPP Research group | Systematic Sampling |
| 21-abr-2020 | Colmenar Viejo | Madrid | 1 | 40.691917 | -3.767194 | 2020 | IVPP Research group | Systematic Sampling |
| 29-abr-2020 | Colmenar Viejo | Madrid | 1 | 40.691917 | -3.767194 | 2020 | IVPP Research group | Systematic Sampling |
| 13-may-2020 | Bustarviejo | Madrid | 3 | 40.691827 | -3.767162 | 2020 | IVPP Research group | Systematic Sampling |
| 29-ene-2021 | Cartagena | Murcia | 1 | 37.611866 | -0.750929 | 2021 | Antonio Montserrat | Systematic sampling |
| 29-ene-2021 | Valle Perdido | Murcia | 1 | 37.927495 | -1.148376 | 2021 | Antonio Montserrat | Systematic sampling |
| 5-feb-2021 | Jumilla | Murcia | 1 | 38.492305 | -1.189841 | 2021 | Antonio Montserrat | Systematic sampling |
| 6-feb-2021 | Jumilla | Murcia | 1 | 38.442847 | -1.316243 | 2021 | Antonio Montserrat | Systematic sampling |
| 17-feb-2021 | Yecla | Murcia | 2 | 38.618762 | -1.127371 | 2021 | Antonio Montserrat | Systematic sampling |
| 17-feb-2021 | Yecla | Murcia | 2 | 38.7307 | -1.13601 | 2021 | Antonio Montserrat | Systematic sampling |
| 19-feb-2021 | Alhama de Murcia | Murcia | 2 | 37.838723 | -1.467839 | 2021 | Antonio Montserrat | Systematic sampling |
| 9-feb-2022 | IMIDRA Alcalá Henares | Madrid | 1 | 40.521133 | -3.290865 | 2022 | Marina Morente | Field observation |
| 13-mar-2018 | Constantina | Sevilla | 2 | 37.862028 | -5.57736391 | 2018 | Jose Manuel Durán | Systematic sampling |
| 19-feb-2021 | El Berro | Murcia | 2 | 37.887945 | -1.493203 | 2021 | Antonio Montserrat | Systematic sampling |
| 18-mar-2018 | Guadalest | Alicante | 3 | 38,662,778 | -0.200278 | 2018 | Marina Morente | Systematic sampling |
| 24-mar-2021 | Jumilla | Murcia | 3 | 38.4447945 | -1.3157153 | 2021 | IVPP Research group | Field observation |
| 24-mar-2021 | Yecla | Murcia | 3 | 38.577469 | -1.1976939 | 2021 | IVPP Research group | Field observation |
| 13-abr-2021 | Sierra Aracena | Huelva | 4 | 37.861016 | -6.484765 | 2021 | IVPP Research group | Field observation |
| 29-abr-2021 | Villanueva Cañada | Madrid | 5 | 40.454418 | -4.005871 | 2021 | IVPP Research group | Field observation |
| 27-may-2021 | Santa María de la Alameda | Madrid | 3 | 40.611108 | -4.263008 | 2021 | IVPP Research group | Field observation |
