## Supporting Document 5 for "Degree-day-based model to predict egg hatching of *Philaenus spumarius* (Hemiptera: Aphrophoridae), the main vector of *Xylella fastidiosa* in Europe"

**Model Calibration considering diapause breakage on January 1^st^**

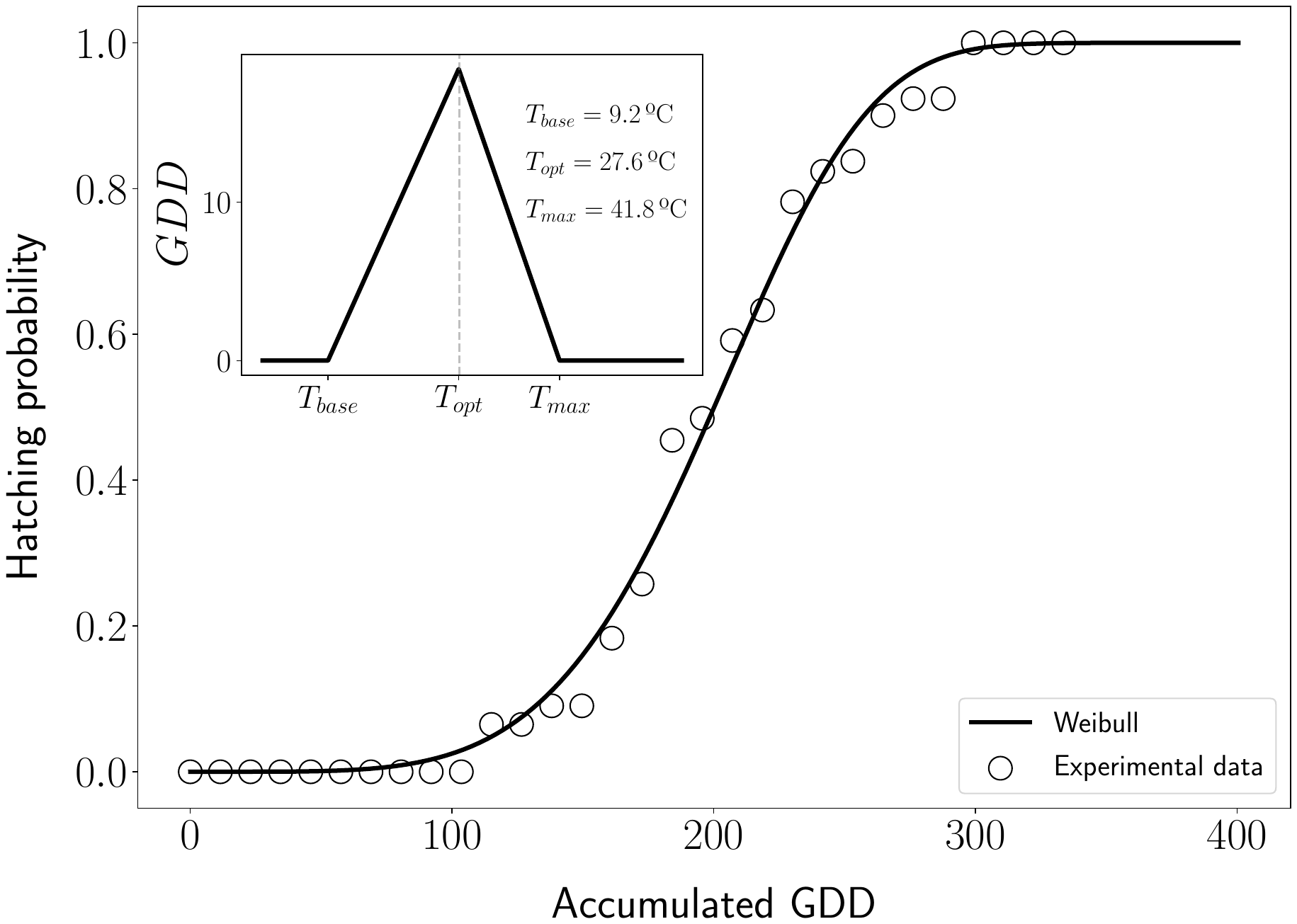

**Model Calibration considering diapause breakage on November 1^st^**

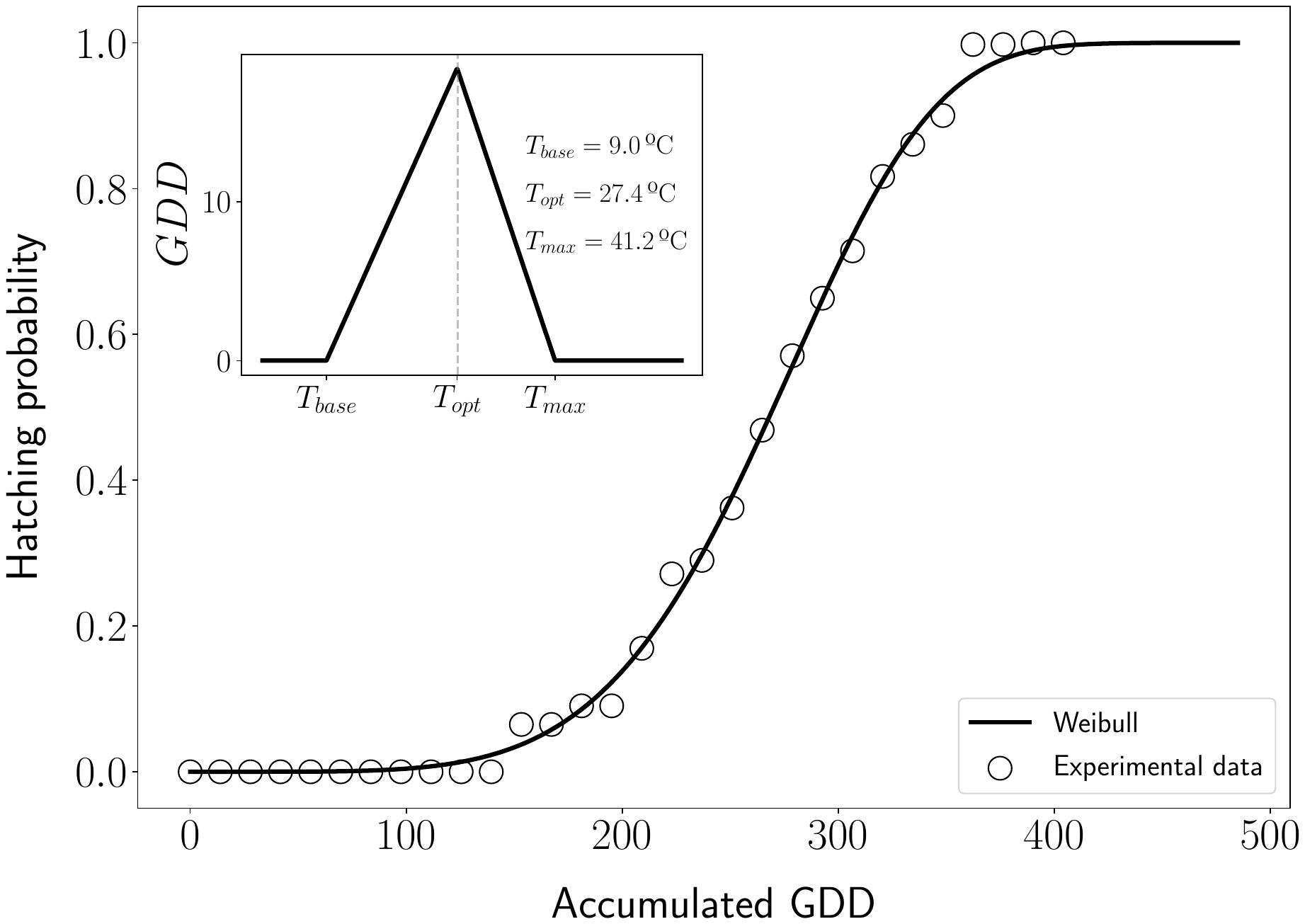
