## Supporting Document S9 for "Degree-day-based model to predict egg hatching of *Philaenus spumarius* (Hemiptera: Aphrophoridae), the main vector of *Xylella fastidiosa* in Europe"

**Document S4.** Monthly mean (± SD) temperatures (°C) and RH (%) measured in the four field sites in Bustarviejo, Mataelpino, Pedrezuela and Alcalá Henares.

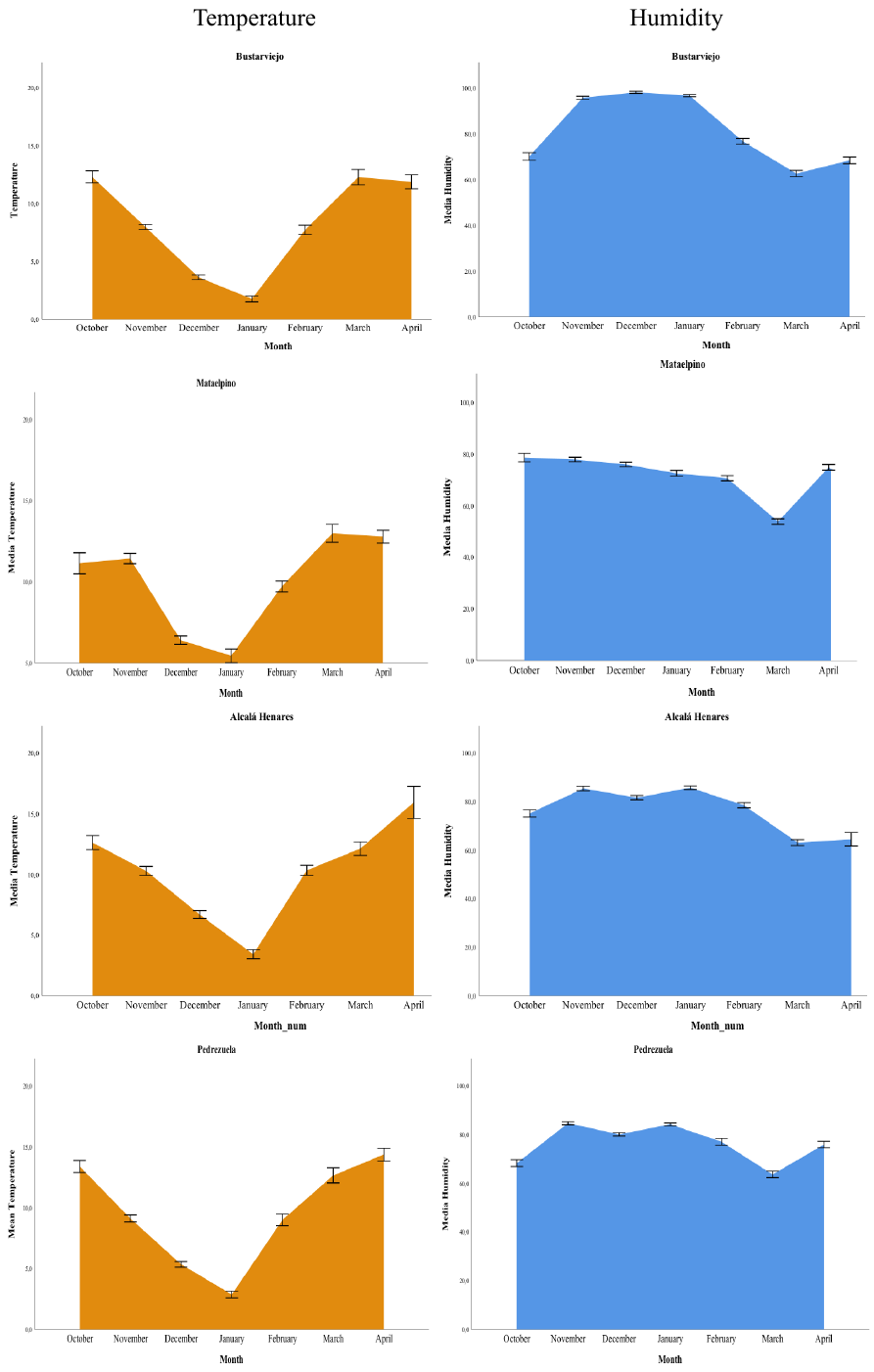
